# Enhancement of alcohol aversion by the nicotinic acetylcholine receptor drug sazetidine-A

**DOI:** 10.1101/723916

**Authors:** Jillienne C. Touchette, Janna K. Moen, Jenna M. Robinson, Anna M. Lee

## Abstract

The prevalence of alcohol use disorders (AUDs) has steadily increased in the United States over the last 30 years. Alcohol acts on multiple receptor systems including the nicotinic acetylcholine receptors (nAChRs), which are known to mediate alcohol consumption and reward. We previously reported that the preclinical drug sazetidine-A, a nAChR agonist, reduces alcohol consumption without affecting nicotine consumption in C57BL/6J mice. Here, we found that sazetidine-A enhances the expression of alcohol aversion without affecting the expression or acquisition of conditioned alcohol reward in C57BL/6J mice. Microinjection of sazetidine-A into the ventral midbrain targeting the ventral tegmental area (VTA) reduced binge alcohol consumption, implicating the neurocircuitries in this region in mediating the effects of sazetidine-A. Furthermore, sazetidine-A-induced reduction in alcohol consumption was mediated by non-α4 containing nAChRs, as sazetidine-A reduced binge alcohol consumption in both α4 knock-out and wild-type mice. Finally, we found that in mice pre-treated with sazetidine-A, alcohol induced *Fos* transcript within *Th*-expressing but not *Gad2*-expressing neurons in the VTA as measured by increased *Fos* transcript expression. In summary, we find that sazetidine-A acts on non-α4 nAChRs to enhance the expression of alcohol aversion, which may underlie the reduction in alcohol consumption induced by sazetidine-A. Elucidating the identity of non-α4 nAChRs in alcohol aversion mechanisms will provide a better understanding the complex role of nAChRs in alcohol addiction and potentially reveal novel drug targets to treat AUDs.

## Introduction

Alcohol use disorders (AUD) have steadily increased in the United States, along with associated increases in high-risk drinking and mortality.^1^ An estimated 30 million Americans met criteria for an AUD in 2013, an increase of 12 million cases compared with the previous decade.^1^ Currently, there are only 3 FDA-approved pharmacotherapies for AUD in the United States (naloxone, acamprosate, disulfiram), all with variable success rates,^2^ which highlights the need to further understand how alcohol interacts with other receptor systems and identify new drug targets.

The neuronal nicotinic acetylcholine receptors (nAChRs) play an important role in mediating the rewarding properties of alcohol. The nAChRs are pentameric cation channels located on presynaptic terminals and cell bodies of neurons, thus modulating cell excitability and neurotransmitter release.^3, 4^ There are 11 nAChR subunits found in the brain (α2-7, α9-10, β2-4) that can combine to form multiple nAChR subtypes, each with its own unique receptor properties such as different ligand binding affinities, calcium permeability and expression patterns.^3^ Alcohol does not directly agonize nAChRs, but modulates nAChR functional activity^5^ and increases the release of acetylcholine (ACh), the endogenous ligand for nAChRs.^6^ Both nAChR antagonists such as mecamylamine (a non-specific antagonist) and partial agonists such as varenicline (an α4β2 partial agonist) reduce alcohol consumption in rodents, illustrating the complex involvement of nAChRs in mediating alcohol consumption.^7–9^ Varenicline has been tested in human trials for alcohol dependence with mixed results, with some studies reporting reduced alcohol consumption^10–12^ but not others.^13, 14^

The preclinical drug sazetidine-A initially agonizes and then desensitizes α4β2, α3β4*, α6* and α7 nAChR subtypes (*denotes additional subunits in the nAChR pentamer).^15–18^ We previously reported an alcohol-specific effect of sazetidine-A in mice; a peripheral injection reduced 24-hour continuous and binge drinking-in-the-dark (DID) alcohol consumption in both male and female C57BL/6J mice without affecting the consumption of water, saccharin, or nicotine.^19^ Here, we investigated the behavioral mechanism by which sazetidine-A reduces alcohol consumption. We report that sazetidine-A enhances the expression of alcohol aversion without affecting the expression or acquisition of alcohol reward. Sazetidine-A does not require the α4 nAChR subunit to reduce alcohol consumption, thus implicating non-α4 containing nAChRs in the mechanism of action of sazetidine-A.

## Materials and Methods

### Animals and Drugs

Eight week old adult male C57BL/6J mice (Jackson Laboratory, Sacramento, CA) acclimated to our facility for a minimum of six days before behavioral experiments. Heterozygous α4 nAChR subunit knock-out (KO) breeder pairs were provided by Dr. Jerry Stitzel at the University of Colorado, Boulder, and α4 nAChR knock-out and their wild-type (WT) littermates were bred on site. All mice were group housed in standard cages under a 12-hour light/dark cycle until the start of behavioral experiments, when they were individually housed. All animal procedures were performed in accordance with the Institutional Animal Care and Use Committee at the University of Minnesota, and conformed to NIH guidelines (National Research Council Committee for the Update of the Guide for the Care and Use of Laboratory Animals, 2010). Alcohol (Decon Labs, King of Prussia, PA) was mixed with tap water for drinking studies or saline to 20% v/v for injections. Sazetidine-A dihydrochloride was purchased from Tocris Bioscience (Bio-techne, Minneapolis, MN), and made fresh for each experiment by dissolving in saline. All peripheral injections were administered intraperitoneally and all experiments were performed in separate groups of drug naïve mice.

### Binge alcohol consumption in female α4 wild-type and knock-out mice

Female mice were used since our previous work showed that sazetidine-A reduces binge consumption in both male and female C57BL/6J mice and female mice consume more alcohol than male.^19^ For the binge drinking-in-the-dark (DID) procedure, mice were habituated to three *i.p.* injections of saline (10μL/g) one week prior to the experiment. Mice were then presented with a bottle of 20% alcohol for 2 hours, starting 2 hours into the dark cycle for 3 consecutive days, based on Rhodes, et al.^20^ The water bottle was removed only when the alcohol bottle was present, but was freely available at all other times. On Day 4, mice were injected with either saline or sazetidine-A (1mg/kg *i.p.*) 1 hour prior to the presentation of the alcohol bottle for 4 hours. Alcohol consumption was calculated by volume, and spillage or evaporation was controlled for by a bottle in an empty control cage.

### Ventral tegmental area (VTA) microinfusion and binge alcohol consumption in male C57BL/6J mice

Male C57BL/6J mice were anesthetized with ketamine/xylazine (80:10 mg/kg *i.p.*) and implanted with bilateral 26 gauge stainless steel guide cannula (Plastics One, Roanoke, VA) targeting the VTA (−3.2mm AP, +/− 0.5mm ML, and −4.96 mm ventral to the surface according to the Paxinos and Franklin mouse brain atlas).^21^ The cannulae were fixed with dental adhesive (Geristore, DenMat, Lompoc, CA). Mice then recovered for 3 weeks in their home cage. One week prior to the binge consumption procedure, mice were habituated to handling and received 3 saline infusions at a volume of 0.5 μL, at a rate of 0.25 μL/min. For the DID procedure, mice were presented with a bottle of 20% alcohol for 2 hours, starting 3 hours into the dark cycle for 4 consecutive days, based on Rhodes, et al.^20^ On Day 5, mice were infused with either saline (1μL) or sazetidine-A (1μg in 1μL) at a rate of 0.25 μL/min, 20 minutes prior to the presentation of the alcohol bottle for 4 hours. Alcohol consumption was calculated by volume, and spillage or evaporation was controlled for by a bottle in an empty control cage. After the completion of the experiment, 1μL of ink was bilaterally injected, brains were removed and sectioned to visualize injection sites. Verification of sites was performed in a blinded manner to the behavioral results. One mouse in the sazetidine-A group and one mouse in the saline group had injections outside of the VTA and were excluded from the analysis.

### Effect of sazetidine-A on the expression and acquisition of alcohol conditioned place preference (CPP)

*Expression:* Male C57BL/6J mice were tested in an unbiased alcohol CPP procedure modified from Cunningham, et al.^22^ The chamber apparatus contained a standard two-compartment place preference insert with different floor textures (Med Associates, St. Albans, VT). The procedure consisted of one habituation session, ten conditioning sessions, one preference test (Test 1), and one test session with sazetidine-A pre-treatment (Test 2). For the habituation session, mice were injected with saline and placed in the apparatus with access to both chambers for 30 minutes, and the baseline time spent in each chamber was recorded. For the conditioning sessions, alcohol (2g/kg *i.p.*) or an equivalent volume of saline was injected and mice were immediately confined to one chamber for 5 minutes. The next day, mice received the opposite injection paired with the alternate chamber, and the pairings were alternated each day for 10 total sessions. Control mice received saline paired with both sides. For the first preference test (Test 1), mice were injected with saline and had access to both sides for 30 minutes. The alcohol CPP index was calculated as the time spent in the alcohol-paired chamber during test day minus time spent in that same chamber on habituation day. The next day, 27 of the 30 mice were randomly divided into 2 groups that were pre-treated with sazetidine-A (1mg/kg, *i.p.*) or saline 1 hour prior to a second 30 min preference test (Test 2). Three mice developed strong aversion to alcohol conditioning (−210 CPA index or lower) and were not randomized for Test 2.

*Acquisition:* To test the effect of sazetidine-A on the acquisition of alcohol CPP, mice underwent a similar procedure described above except that mice were pre-treated with sazetidine-A (1mg/kg, *i.p.*) or saline, 1 hour before the alcohol-paired session. For the saline-paired sessions, mice received a pre-treatment saline injection. The alcohol CPP index was calculated as the time spent in the alcohol-paired chamber during test day minus time spent in that same chamber on habituation day.

### Effect of sazetidine-A and mecamylamine on the expression of alcohol conditioned place aversion (CPA)

Male mice were tested in an unbiased alcohol CPA procedure modified from Cunningham, et al., ^23^ which was similar to the alcohol CPP procedure except that the mice received an injection of alcohol (2g/kg, *i.p.*) or saline immediately upon removal from the chamber, prior to being placed back in their homecage. Control mice received saline paired with both sides. For the first aversion test (Test 1), mice were injected with saline and had access to both sides for 30 minutes. The alcohol CPA index was calculated as the time spent in the alcohol-paired chamber during test day minus time spent in that same chamber on habituation day. Only the alcohol-conditioned mice that had a negative CPA index (22 out of 30 mice) proceeded to be randomized into 2 groups that were pre-treated with sazetidine-A (1mg/kg, *i.p.*) or saline 1 hour prior to a second 30 min aversion test (Test 2).

To test the effects of mecamylamine on the expression of alcohol aversion, a separate alcohol CPA was conducted in a different cohort of mice. All mice were conditioned for alcohol aversion with 2 g/kg alcohol as described above. All mice expressed a negative CPA index on Test day 1, thus the next day all mice were randomized into 3 groups and pre-treated with saline or mecamylamine (2 or 3mg/kg *i.p.*) 20 minutes prior to the second aversion test (Test 2). The alcohol CPA index was calculated as the time spent in the alcohol-paired chamber during test day minus time spent in that same chamber on habituation day.

### Place conditioning with sazetidine-A

Male C57BL/6 mice were tested in a place conditioning procedure consisting of one habituation session, ten daily conditioning sessions and one test. For the habituation session, mice were injected with saline and had access to both chambers for 30 minutes. For the conditioning sessions, mice were treated with sazetidine-A (1mg/kg *i.p.*) or saline 1 hour prior to the 30-minute conditioning session. The next day, mice received the opposite injection paired with the alternate chamber, and the pairings were alternated each day. Control mice received saline paired with both sides. For the test session, mice were injected with saline and had access to both chambers for 30 minutes. The conditioning index was calculated similar to the alcohol CPP index.

### Alcohol loss-of-righting reflex (LORR)

Male C57BL/6J mice were pre-treated for 1 hour with sazetidine-A (1mg/kg, *i.p.*) or saline. Mice were then injected with 4g/kg alcohol *i.p.* and placed on their backs, alone, in clean cages. Righting reflex was considered lost when the mouse failed to right itself for at least 30 seconds after administration of the alcohol injection. The mouse was considered to have recovered the righting reflex when it could right itself three times within 30 seconds.

### Locomotor activity

Male C57BL/6J mice were pre-treated with sazetidine (1mg/kg, *i.p.*) or saline for 1 hour and placed back in their home cage. The mice were then placed in a locomotor chamber (Med Associates, St. Albans, VT) for 1 hour. Total distance traveled, measured via infrared beam breaks, was measured in 5 minute time blocks and compared across the two treatment groups.

### Fluorescent in situ hybridization (FISH)

To compare the effect of alcohol and sazetidine-A on *Fos* transcript expression within neuronal subpopulations, mice were pre-treated with saline for 1 hour, placed back in their home cage, and then injected with 1) saline (saline-saline group), 2) alcohol (saline-alcohol group) or 3) sazetidine-A (saline-SAZ group). A fourth group was pre-treated with 1mg/kg sazetidine-A for 1 hour and then injected with 2g/kg alcohol (SAZ-alcohol group). Mice were then placed back in their home cage for 90 minutes after the second injection. Mice were then sacrificed and brains were removed, snap frozen in isopentane, and sectioned on a cryostat (HM 525 NX, ThermoFischer) into 16 μM sections. Sections were chosen based on stereotaxic landmarks (acceptance criteria: −2.8 and −3.2 relative to bregma based on the Paxinos and Franklin mouse brain atlas).^21^ Separate sections were collected and processed for co-localization of *Fos*+*Th*, and *Fos*+*Gad2*. The *Th* and *Gad* sections were collected consecutively to provide anatomical matching. Sections were adhered to Superfrost® Plus slides, kept at −20°C for 60 minutes to dry and stored at −80°C until use. Sections were fixed with 4% PFA for 1 hour and processed for RNAScope (Advanced Cell Diagnostics) multichannel FISH according to manufacturer instructions for fluorescent multiplex assays. Sections were counterstained with DAPI for 20 seconds at room temperature, coverslipped with Prolong Gold Antifade (ThermoFisher Scientific), and stored at 4°C. Probes for detection of specific targets *(Fos, Th*, *Gad2*) were purchased from Advanced Cell Diagnostics (ACD; http://acdbio.com/).

### FISH image acquisition and analysis

Sections containing the VTA were imaged on a Keyence BZ-X700 epifluorescent microscope at 20X magnification. Images for each channel were obtained in z-stack and stitched using Keyence analysis software, and all images were acquired and processed in the same manner. Multichannel images were opened and all channels (including DAPI) were overlaid. A fixed area was imposed around the VTA based on stereotaxic landmarks to establish regional boundaries. Cells within the boundaries of the VTA were considered positive for each transcript when at least 5 puncta were observed within each cell, and cell counts were tracked by a blinded researcher using the Cell Counter plugin in ImageJ v1.52e.^24^ All counts were made bilaterally, with 3-4 mice analyzed for each treatment group, with 4 slices (2 *TH+Fos* and 2 *GAD+Fos*) per mouse.

### Statistical Analysis

All analyses were calculated using Prism 8.0 (GraphPad, La Jolla, CA). Data were tested for normality and variance, and outliers were detected using the Grubb’s test. Welch’s corrections were used if variances were unequal. Comparison of data across time used two-way repeated measures ANOVA followed by Tukey’s multiple comparisons tests (ex. α4 nAChR DID, locomotor activity). Comparisons between groups with a single dependent variable used Student’s *t*-tests or one-way ANOVAs followed by multiple comparisons tests (ex. FISH RNAScope).

For the conditioning data, comparison across treatment groups was analyzed using Student’s *t*-tests or one-way ANOVA followed by multiple comparisons test (Dunnett’s multiple comparison tests or unpaired *t*-tests with Welch’s corrections). To assess the development of conditioning within each group, we used a one-sample t-tests compared with a conditioning index of zero (no reward or aversion).

## Results

### Sazetidine-A reduced binge alcohol consumption inα4 knock-out mice

We previously showed that systemic sazetidine-A reduced binge DID alcohol consumption in both male and female C57BL/6J mice.^19^ Since a major nAChR subtype that is activated and desensitized by sazetidine-A is the α4β2 nAChR, we sought to determine whether the α4 nAChR subunit was important for the actions of sazetidine-A. We tested the effect of systemic sazetidine-A in mice that lack the α4 nAChR subunit in the DID procedure. We found that sazetidine-A (1mg/kg *i.p.*) reduced alcohol consumption in both α4 WT and KO female mice, as there was a main effect of drug treatment with no effect of genotype or an interaction between drug treatment and genotype (**Fig. 1**, 2-way ANOVA F_interaction_(1,27)=0.281, *P*=0.60; F_treatment_(1,27)=13.08, *P*=0.001; F_genotype_(1,27)=0.004, *P*=0.95). Thus, these data suggest that sazetidine-A reduces alcohol consumption by acting on non-α4* nAChRs.

**Fig. 1.**
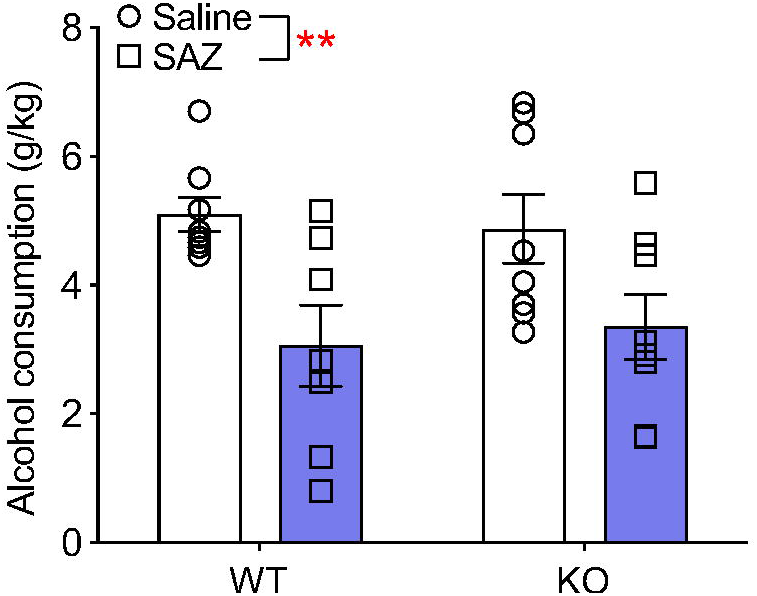
Sazetidine-A reduces binge alcohol consumption in α4 knock-out mice. **(A)**Sazetidine-A (1 mg/kg *i.p.*) reduced binge alcohol consumption in both wild-type and α4 knock-out female mice. *n*=7-8 per group. 2-way ANOVA for drug treatment: F_treatment_(1,27)=13.08, \*\**P*=0.001.

### Microinjection of sazetidine-A into the ventral midbrain reduced alcohol consumption

As our prior work used systemic injections of sazetidine-A, we tested whether administration of sazetidine-A into the ventral midbrain targeting the VTA would reduce acute binge DID alcohol consumption. Male C57BL/6J mice were bilaterally implanted with cannula targeting the VTA prior to a binge DID procedure. Mice microinjected with 1μg/μL sazetidine-A consumed significantly less alcohol during the 4h binge session compared with mice infused with saline (**Fig. 2**, unpaired t-test, *t*=2.965, df=9, *P*=0.02).

**Fig. 2.**
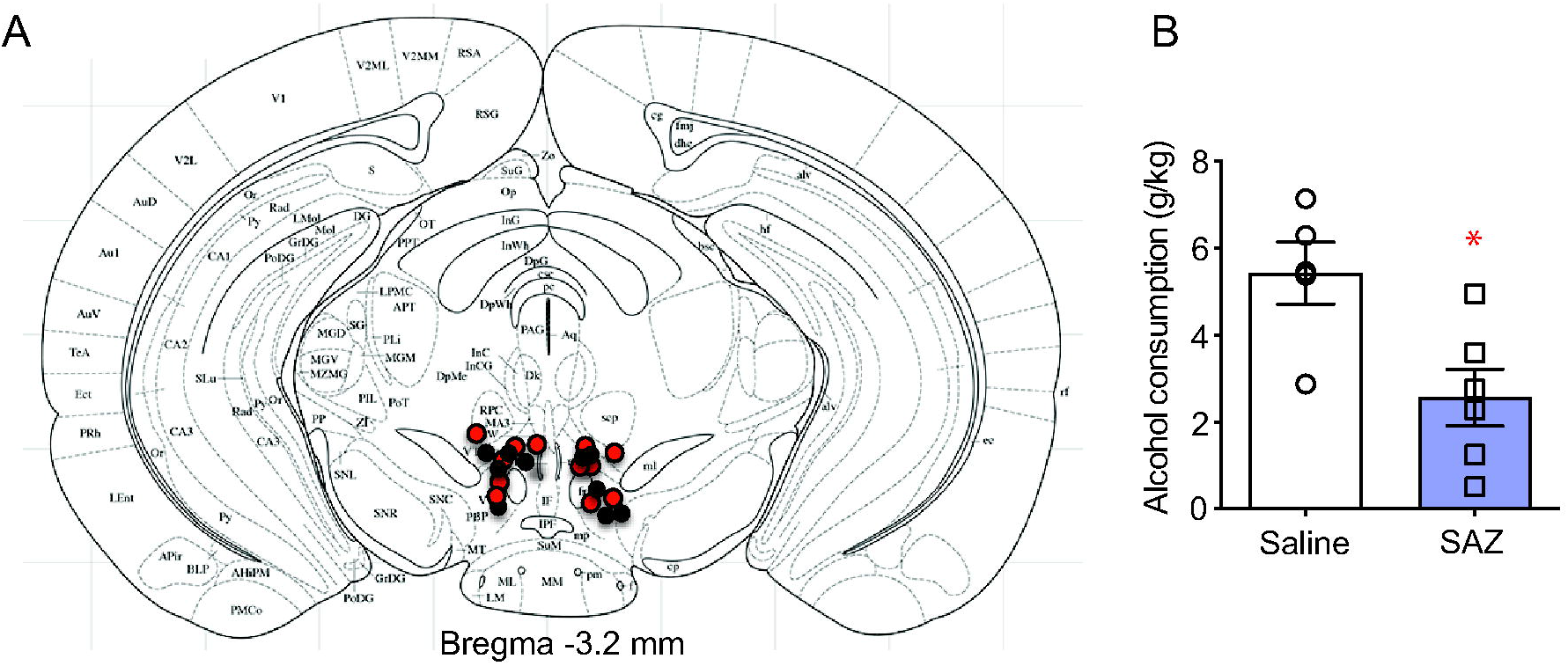
Microinfusion of sazetidine-A into the ventral midbrain targeting the VTA reduces binge alcohol consumption. **(A)**Coronal mouse brain atlas diagrams from mouse brain atlas showing the confirmed injection sites in mice infused with sazetidine-A (1 μg/μL, red) or saline (1 μL, black). **(B)**Average alcohol consumption on binge day in male C57BL/6J mice pre-treated with saline or sazetidine-A 20 min prior to alcohol access. \**P*=0.02, *n*=5-6 per group, mean±SEM.

### Sazetidine-A enhanced the expression of alcohol aversion without affecting alcohol reward

To determine the behavioral mechanism by which a single injection of sazetidine-A reduced DID alcohol consumption, we first tested the effect of sazetidine-A on the expression of alcohol reward using conditioned place preference (CPP). Male C57BL/6J mice were conditioned with 2g/kg *i.p.* alcohol injections, which produced significant preference for the alcohol-paired chamber compared with saline conditioning on preference test 1 (**Fig. 3A**, Student’s *t*-test between saline and alcohol groups: *t*=2.675, df=39, *P*=0.01; one-sample *t*-test compared with a CPP index of zero: saline group *t*=1.384, df=10, *P*=0.20; alcohol group *t*=3.123, df=29, *P*=0.004). The next day, the alcohol-conditioned mice were randomized into 2 groups that were pre-treated with sazetidine-A (1mg/kg *i.p.*) or saline 1 hour prior to a second alcohol preference test. We found no significant difference in the alcohol CPP indices indicating that sazetidine-A does not impact the expression of alcohol CPP (**Fig. 3B**, *t*=0.191, df=25, *P*=0.85).

**Fig. 3.**
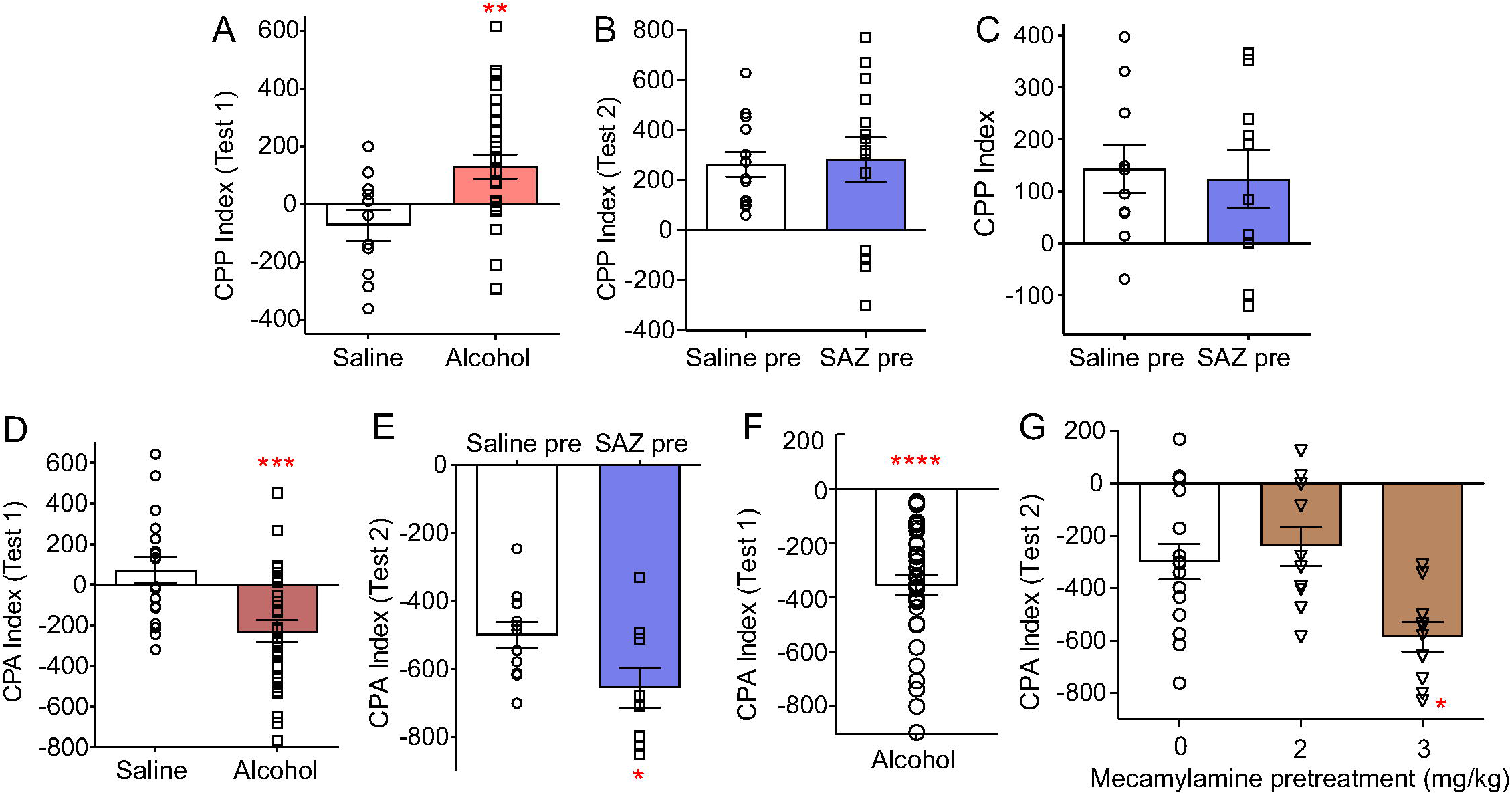
Sazetidine-A enhanced expression of alcohol aversion without affecting the expression or acquisition of alcohol reward. **(A)**Conditioning with 2 g/kg alcohol resulted in preference for the alcohol-paired chamber (Test 1), whereas conditioning with saline produced no significant preference. \*\**P*<0.05 using a one-sample *t*-test compared with a CPP index of zero, *n*=30 alcohol, *n*=11 saline. **(B)**Pre-treatment with sazetidine-A (1 mg/kg *i.p.*) 1h prior to the second test day (Test 2) did not produce significant differences in alcohol CPP versus saline pre-treated mice, *n*=13-14 per group. **(C)**There was no difference in the CPP index between mice pre-treated with sazetidine-A (1 mg/kg *i.p.*) or saline prior to each alcohol conditioning session, *n*=10 per group. **(D)**Aversion conditioning with 2 g/kg alcohol resulted in aversion to the alcohol-paired chamber (Test 1), whereas conditioning with saline produced no significant aversion. \*\*\**P*=0.0001 using one-sample *t*-test compared with a CPA index of zero, *n*=30 alcohol, *n*=18 saline. **(E)**Pre-treatment with sazetidine-A (1 mg/kg *i.p.*) 1h prior to the second test day (Test 2) enhanced alcohol aversion. \**P*=0.03 compared to the saline group, *n*=9-11 per group. **(F)**Aversion conditioning with 2 g/kg alcohol produced significant alcohol CPA. \*\*\*\**P*<0.0001 using one-sample t-test compared with a CPA index of zero, *n*=35. **(G)**Pre-treatment with mecamylamine at 3 mg/kg *i.p.* 20 min prior to the second test day (Test 2) enhanced alcohol aversion. \**P*<0.05 compared to saline group by Dunnett’s post-hoc test, *n*=10-15 per group. All conditioning indices are in minutes, all mice used were male C57BL/6J, all data shown as mean±SEM.

We then tested whether sazetidine-A impaired the acquisition of alcohol reward during the conditioning procedure with a separate group of mice. Sazetidine-A (1mg/kg) or saline was injected 1 hour prior to each alcohol conditioning session. We found no significant difference in alcohol CPP index on test day between the sazetidine-A and saline pre-treated groups, indicating that sazetidine-A does not impact the acquisition of alcohol CPP (**Fig. 3C**, *t*=0.262, df=18, *P*=0.79; one-sample *t*-test compared with a CPP index of zero: saline pre-treated group: *t*=3.106, df=9, *P*=0.01; sazetidine-A pre-treated group: *t*=2.250, df=9, *P*=0.05).

We then tested whether sazetidine-A affected the expression of alcohol aversion using conditioned place aversion (CPA). Mice were conditioned with 2g/kg alcohol injections, which produced significant aversion for the alcohol-paired chamber compared with saline conditioning (**Fig. 3D**, Student’s *t*-tests between saline and alcohol-treated groups *t*=4.204, df=45, *P*=0.0001; one-sample *t*-test compared with a CPA index of zero: saline-treated group *t*=1.18, df=17, *P*=0.25; alcohol-treated group *t*=4.47, df=28, *P*=0.001). The next day, the alcohol-conditioned mice that developed aversion were randomized into 2 groups that were pre-treated with sazetidine-A (1mg/kg) or saline prior to a second aversion test. We found that pre-treatment with sazetidine-A enhanced the expression of alcohol CPA compared with saline (**Fig. 3E**, *t*=2.284, df=18, *P*=0.03).

Since sazetidine-A initially activates and then desensitizes nAChRs, we sought to determine whether nAChR activation or desensitization was important for the enhancement of alcohol aversion expression. We tested the effect of mecamylamine, a non-specific nAChR antagonist, on the expression of alcohol aversion in a separate group of mice. Alcohol aversion conditioning produced significant aversion (**Fig. 3F**, one-sample *t*-test compared with a CPA index of zero: *t*=9.655, df=34, *P*<0.0001). The next day, the alcohol-conditioned mice were randomized into 3 groups and were pre-treated with mecamylamine (0, 2 or 3 mg/kg) 20 minutes prior to a second aversion test. Mecamylamine enhanced the expression of alcohol aversion at the 3 mg/kg dose but not the 2 mg/kg dose, compared with the saline pre-treated group (**Fig. 3G**, one-way ANOVA F(2,32)=6.38, *P*=0.005). These results suggest that antagonism of nAChRs, rather than activation of nAChRs, mediates the enhanced expression of alcohol aversion.

### Sazetidine-A enhanced the sedative-hypnotic effects of alcohol

To determine whether sazetidine-A affected the sedative-hypnotic effects of high-dose alcohol, we tested whether sazetidine-A altered acute alcohol-induced loss-of-righting-reflex (LORR). Pre-treatment with sazetidine-A (1mg/kg) for 1 hour enhanced alcohol-induced LORR duration produced by 4g/kg alcohol compared with saline pre-treatment (**Fig. 4A**, *t*=2.554, df=7.477, *P*=0.04).

**Fig. 4.**
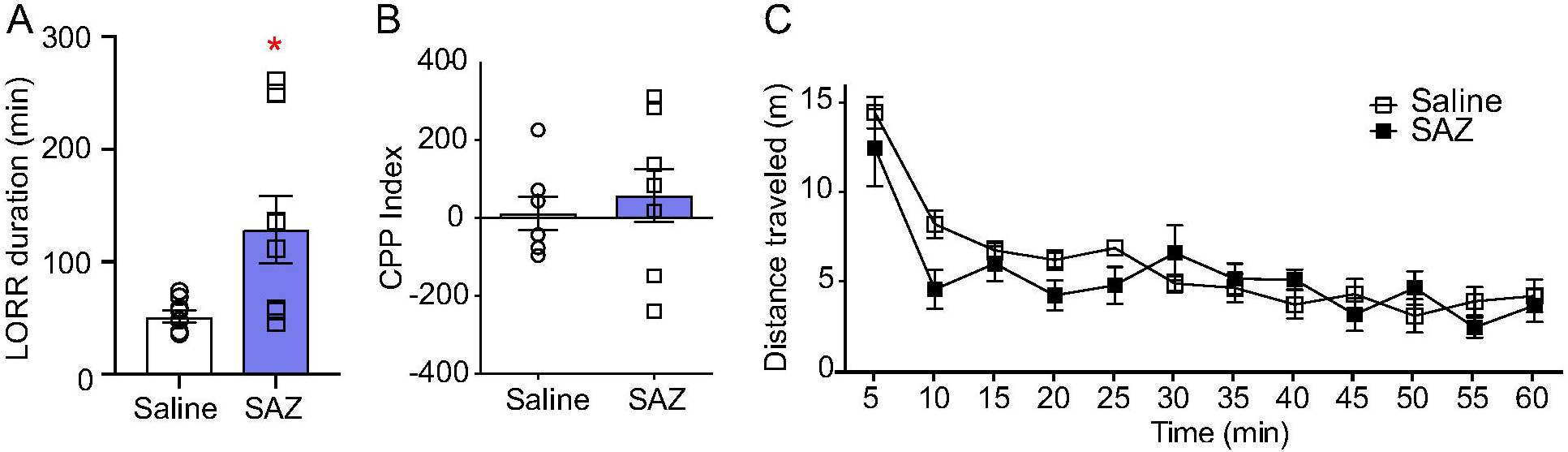
Effect of sazetidine-A on alcohol-induced LORR, place conditioning and locomotor activity. **(A)**Average alcohol-induced LORR duration from 4g/kg *i.p.* alcohol in mice pre-treated with sazetidine-A (1 mg/kg, *i.p.*) was greater than mice pre-treated with saline. \**P*=0.04 compared with saline treatment, *n*=7-8 per group. **(B)**Place conditioning with sazetidine-A only using a 1mg/kg *i.p.* dose and 1h timing was not significantly different from saline conditioning, *n*=7-8 per group. **(C)**Average ambulatory distance in mice after injection with sazetidine-A (1mg/kg, *i.p.*, 1h) was not significantly different from the saline treated group, *n*=8 per group. All data shown as mean±SEM.

### Sazetidine-A treatment does not affect place conditioning or locomotor activity

All our alcohol consumption, conditioning experiments and LORR treatments used a 1 hour pre-treatment of 1mg/kg sazetidine-A. We tested whether this dose and timing of sazetidine-A had any rewarding or aversive effects alone in place conditioning, and whether it affected locomotor activity. We found that 1mg/kg sazetidine-A produced no expression of reward or aversion, and was similar to the conditioning produced by saline (**Fig. 4B**, *t*=0.559, df=13, *P*=0.59). We also found no main effect of sazetidine-A treatment on locomotor activity as measured by beam breaks (**Fig. 4C**, 2-way RM ANOVA F_interaction_(11,154)=4.75, *P*=0.07; F_treatment_(1,154)=0.89, *P*=0.22; F_time_(11,154)=48.80, *P*<0.0001).

### Sazetidine-A induced neuronal activation in the VTA primarily in TH-expressing neurons

As a single systemic injection of sazetidine-A reduced DID alcohol consumption and enhanced the expression of alcohol aversion, and micro-injecting sazetidine-A into the ventral midbrain targeting the VTA reduced binge DID consumption, we sought to determine the effect of a single, systemic injection of sazetidine-A on neuronal activity in within the VTA. We determined whether the sazetidine-A induced c-Fos expression occurred in dopamine (DA) or γ-aminobutyric acid (GABA) neurons in the VTA. Fluorescence *in situ* hybridization (RNAScope) was used to identify *Fos* transcript expression in VTA cells that express tyrosine hydroxylase (*Th*), a common transcript in DA neurons, or glutamate decarboxylase 2 (*Gad2*), a transcript expressed in GABA neurons **(Fig. 5A)**. Pre-treatment with sazetidine-A (1mg/kg) prior to a 2g/kg alcohol injection produced an increase in *Fos* expression in *Th-*, but not *Gad2*-expressing neurons, compared with saline pre-treatment (one-way ANOVA F(3,11)=19.53, *P*=0.0001; Tukey’s multiple comparisons test \*\*\**P*<0.001 for SAZ/alc compared to all other groups, **Fig. 5B-C**). There was no significant increase in the number of *Fos*-expressing *Th*-positive cells between the saline, alcohol and sazetidine-A only treated groups, suggesting that the increase in *Fos* expression in *Th-* expressing neurons is not due to either sazetidine-A or alcohol alone (Tukey’s multiple comparisons tests, *P*>0.05 between sal/sal, sal/alc and sal/SAZ). *Fos* expression in *Gad2*-expressing neurons did not change across treatment conditions (one-way ANOVA F(3,11)=1.356, *P*=0.31, **Fig. 5D-E**). These data suggest that pre-treatment with sazetidine-A followed by alcohol injection selectively increased *Fos* expression in *Th*- expressing neurons of the VTA.

**Fig. 5.**
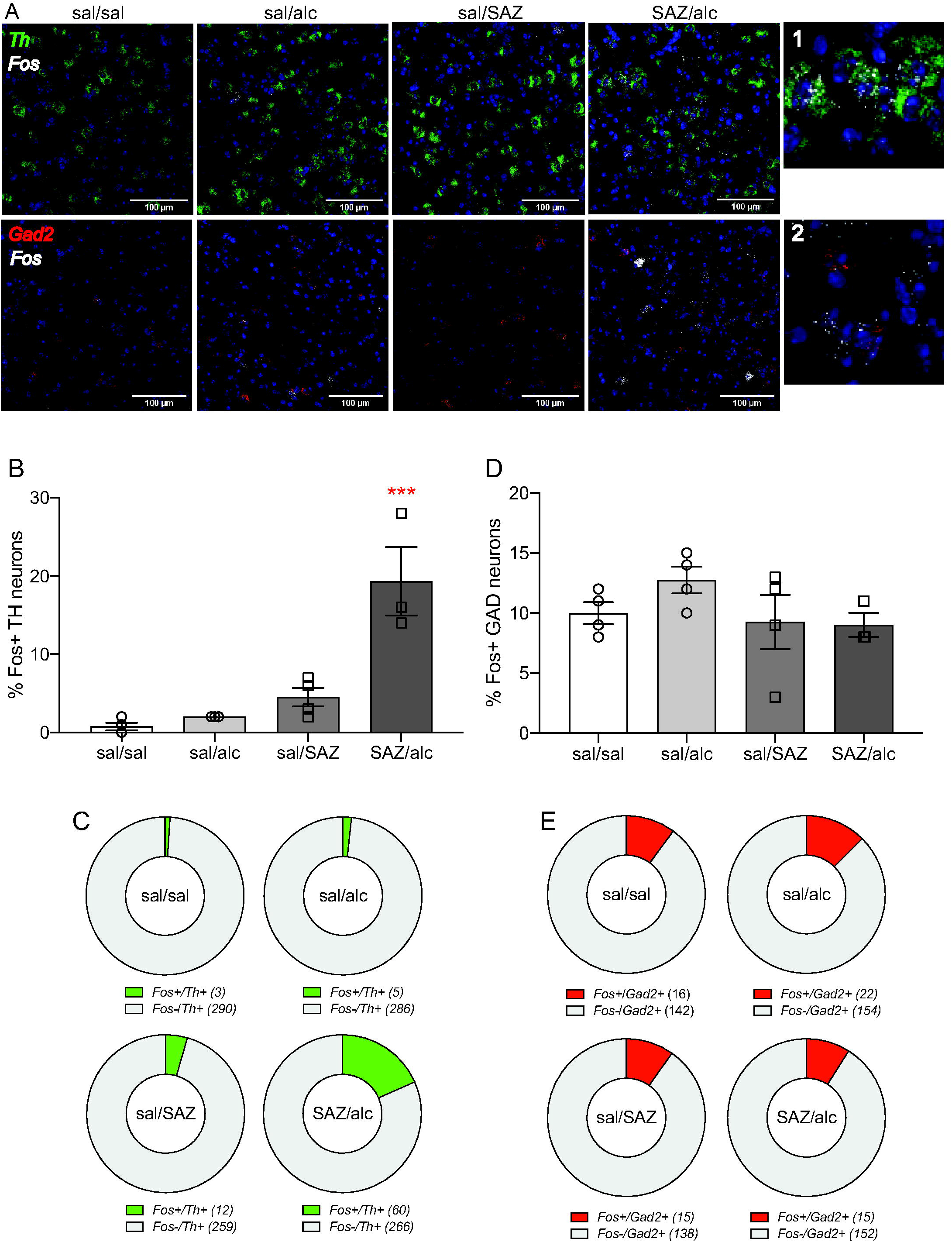
Sazetidine-A pre-treatment followed by alcohol injection induced Fos expression in Th-expressing VTA neurons. **(A)**Representative images of *Fos* (white, Cy5) expression in *Th*- (green, mCherry) or*Gad2*-positive neurons (red, mCherry) in the VTA. Scale bar = 100 μm. *1,2*: magnification of *Th* and *Gad2* in the SAZ/alc group. **(B-C)**The number of *Fos*-expressing *Th-*positive neurons was increased in the VTA of alcohol-injected mice pre-treated with sazetidine-A compared with saline pre-treatment (\*\*\**P*<0.001 for SAZ/alc versus sal/alc), and compared with groups that received only saline or sazetidine-A (\*\*\**P*<0.001 for SAZ/alc versus sal/sal and sal/SAZ). **(D-E)***Fos*-expressing *Gad2*-positive neurons was similar between all treatment groups. *n*=3-4 mice per group, mean ± SEM for **B, D**and averaged group data for **C, E**.

## Discussion

Pharmacological manipulation of nAChRs has been shown to reduce alcohol consumption in animal models, as mecamylamine, varenicline and sazetidine-A reduce alcohol consumption in rodents despite their different mechanisms of actions and nAChR subtype targets.^7–9, 19^ Varenicline is best known as a partial agonist at α4β2 receptors, but *in vitro* studies show that it is also a partial agonist at α3* and α6* receptors and a full agonist at α7 receptors.^16, 25–27^ *In vitro* studies show that sazetidine-A is an agonist at α4β2, α3β4*, α6* and α7 nAChRs,^15–18^ thus sazetidine-A and varenicline act on similar nAChR subtypes but with differing potencies.^15^ While varenicline requires the α4 nAChR subunit to reduce alcohol consumption,^28^ we found that sazetidine-A decreased binge alcohol consumption in both α4 KO and WT mice, indicating that sazetidine-A does not require the α4 subunit to decrease alcohol consumption. A study by Turner and colleagues also demonstrated dissociable effects of varenicline and sazetidine-A in nicotine withdrawal signs in mice.^29^ Furthermore, acute varenicline injection, but not acute sazetidine-A injection, produces anxiolytic effects in the marble-burying test and in novelty-induced hypophagia in mice.^30^ Sazetidine-A, but not varenicline, produced anti-depressant effects on the forced swim test and tail suspension test in mice.^30^ Together, these data suggest that despite the similarity in nAChR targets *in vitro*, sazetidine-A and varenicline likely act through different nAChR subtypes *in vivo* to produce distinct behavioral effects.

A decrease in alcohol consumption may be due to a decrease in alcohol reward and/or an increase in alcohol aversion. We found no effect of sazetidine-A on the expression or acquisition of alcohol CPP in male C57BL/6J mice. In contrast, the non-specific nAChR antagonist mecamylamine has been shown to inhibit the acquisition and expression of alcohol CPP in mice when administered intracerebroventricularly,^31^ which suggests that antagonism of nAChRs reduces the development alcohol reward associations and the expression of such previously learned associations. Interestingly, varenicline has been shown to have no effect on the expression of alcohol CPP in mice,^32^ similar to our findings with sazetidine-A. Notably, the role of nAChRs in alcohol aversion has not been previously explored in pre-clinical models. Here, we found that 1mg/kg sazetidine-A and 3mg/kg mecamylamine enhanced the expression of conditioned alcohol aversion. As sazetidine-A initially activates and then desensitizes nAChRs,^15–18^ and we showed that both sazetidine-A and mecamylamine enhance alcohol aversion, we speculate that reduced nAChR function, either through receptor desensitization or antagonism, mediates the enhanced expression of alcohol aversion. A 4mg/kg mecamylamine dose does not affect locomotor activity or produce aberrant behavior in mice,^33^ thus we believe the lower 3mg/kg used in our study would not produce changes in locomotor activity that would affect the expression of CPA alone.

Overall, these data demonstrate an important role for nAChRs in conditioned alcohol aversion that had not been previously demonstrated, and suggest that enhancing alcohol aversion can be one mechanism by which nAChR drugs mediate a decrease in alcohol consumption. Interestingly, the opiate antagonist naloxone, a clinically approved medication for alcohol dependence, has also been shown to have no effect on expression of alcohol CPP, yet enhances expression of alcohol CPA.^34^ Therefore, developing additional pharmacological agents that increase the expression of previously conditioned alcohol aversion may be a potentially useful and viable strategy to reduce alcohol consumption.

We microinjected sazetidine-A injected directly into the ventral midbrain targeting the VTA and found that it reduced binge alcohol consumption in C57BL/6 mice. We targeted the VTA as it is an important structure that is critical for drug consumption, reward and aversion, and expresses 8 of the 11 nAChR subunits found in the human brain.^35, 36^ The VTA is a heterogeneous structure and contains DA, GABA and glutamate neurons which form distinct circuits that mediate reward and aversion.^37, 38^ In particular, optogenetic activation of VTA GABA neurons or inhibition of VTA DA neurons during place conditioning has been shown to produce conditioned place aversion.^39^ To begin identifying which circuits in the VTA were important for sazetidine-A’s effects, we determined which neuronal cell types within the VTA were activated by injections of 1mg/kg sazetidine-A, 2g/kg alcohol, or an injection of sazetidine-A followed by an injection of alcohol. Examination of transcript expression using FISH RNAScope showed that pre-treatment with 1mg/kg sazetidine-A followed by 2g/kg alcohol (SAZ-alc group) increased *Fos* transcript expression in *Th-*expressing neurons, compared with the saline-alc group. There was no significant increase in *Fos* transcript expression in *Th-* expressing neurons in the saline or sazetidine-A only groups. Moreover, there was no change in the number of *Gad2-*expressing neurons that showed *Fos* expression across all groups, suggesting that GABA neurons in the VTA, whether interneurons or projections neurons, are not activated by SAZ-alcohol injections.

Based on our findings and the known actions of sazetidine-A *in vitro,*^15–18^ we speculate that sazetidine-A agonism of nAChRs produces neuronal activation in *Th*- expressing cells, which then leads to the induction of *Fos* expression. This would be followed by prolonged sazetidine-A induced nAChR desensitization, as sazetidine-A is cleared slowly from the brain and can occupy brain nAChRs for ~8 hours.^40^ Since nAChR desensitization is thought to decrease neuronal activity,^41, 42^ sazetidine-A-induced desensitization may lead to decreased activity of putative VTA DA neurons, thus enhancing alcohol aversion and reducing alcohol consumption. Since we showed that sazetidine-A did not require the α4 nAChR subunit to reduce alcohol consumption, our data suggest that non-α4 nAChRs on *Th*-expressing neurons may important in sazetidine-A mediated alcohol aversion. Future studies will test this working hypothesis and determine which nAChR subunits are expressed in *Fos* expressing *Th*-positive neurons.

A limitation of our micro-injection study is the 1μL volume injected per side, which likely covered a large region that included the VTA. Thus, other brain regions in close proximity to the VTA, such as the substantia nigra and rostromedial tegmental area, may also be contributing to sazetidine-A’s effects. Identifying whether these brain structures are involved in sazetidine-A’s actions on alcohol consumption and aversion will provide further insight into the role of nAChRs in the neurobiology of alcohol’s effects.

## Implications

In summary, we found that sazetidine-A, a nAChR agonist and desensitizer, enhanced the expression of alcohol conditioned place aversion without affecting the expression or acquisition of alcohol conditioned place preference. The enhanced alcohol aversion may be mediated by desensitization or antagonism of nAChRs, as mecamylamine, a nAChR antagonist, also enhanced the expression of alcohol conditioned place aversion. Lastly, our data suggest that the combination of sazetidine-A and alcohol activates dopamine neurons in the VTA, as *Fos* transcript expression was increased in *Th*-, and not *Gad2-*, expressing neurons in the VTA. We speculate that the increase in alcohol aversion contribute to the reduction in alcohol consumption we observed after treatment with sazetidine-A. Further elucidating the role of specific nAChR subunits in alcohol aversion mechanisms may identify novel drug targets for the development of pharmacotherapies to treat alcohol use disorder.

## Acknowledgements

We would like to thank Drs. Julia Lemos and Jeff Stolley for their assistance with the RNA Scope assay, and Dr. Jerry Stitzel for providing the α4 nAChR subunit knock-out breeder pairs. We also acknowledge Jamie Maertens and Cecilia Huffman for their technical assistance in this study.

